# A zebrafish model for *Mycobacterium leprae* granulomatous infection

**DOI:** 10.1101/127639

**Authors:** Cressida A. Madigan, James Cameron, Lalita Ramakrishnan

**Author notes:** Present address: Division of Dermatology, Department of Medicine, David Geffen School of Medicine, University of California, Los Angeles, CA, USA, 90095. Present address: Environmental and Fisheries Science Division, National Oceanic and Atmospheric Administration, Seattle, WA 98115, USA.

## Abstract

Understanding the pathogenesis of leprosy granulomas has been hindered by a paucity of tractable experimental animal models. *Mycobacterium leprae*, which causes leprosy, grows optimally at ~30°C, so we sought to model granulomatous disease in the ectothermic zebrafish. We find noncaseating granulomas develop rapidly, and eventually eradicate infection. *rag1* mutant zebrafish, which lack lymphocytes, also form noncaseating granulomas with similar kinetics, but these control infection more slowly. Our findings establish the zebrafish as a facile, genetically tractable model for leprosy, and reveal the interplay between innate and adaptive immune determinants mediating leprosy granuloma formation and function.

## Introduction

Few animal models exist for the study of *M. leprae* pathogenesis *in vivo*, largely because the ≥37°C core temperature of traditional rodent models prevents *M. leprae* survival [1]. *M. leprae* is propagated for research use in the athymic mouse footpad [1], where they induce granuloma formation but not the neurological disease typical of human leprosy [2]. Armadillos develop neurological disease and form granulomas in response to *M. leprae;* however, they do not breed in captivity and lack most genetic, molecular and immunological tools [3]. Cultured macrophages have been used to model early granuloma formation with *M. leprae*, but the scope of this model remains limited [4]. Overall, the host determinants that mediate granuloma formation in leprosy and their role in pathogenesis are incompletely understood.

The zebrafish has become an effective model for studying *Mycobacterium tuberculosis* granulomas using *M. marinum*, the agent of fish tuberculosis, and a close genetic relative of the *M. tuberculosis* complex [5]. *M. marinum* infection of adult zebrafish results in organized, multicentric granulomas that become necrotic, similar to those of human tuberculosis [6]. Zebrafish are housed at ~30°C, similar to the growth optimum of *M. leprae*; indeed, a more than century-old paper reports experimental *M. leprae* infection of several fish species [7]. Therefore, we explored the zebrafish as a leprosy model, with a focus on granuloma development, fate and function.

## Methods

Zebrafish husbandry and experiments were conducted at the University of Washington in compliance with guidelines from the U.S. National Institutes of Health and approved by the University of Washington Institutional Animal Care and Use Committee. Four-month old male zebrafish, either wildtype AB strain, or sibling *rag1*^*t*26683/*t*26683^ mutants and *rag1*^+/*t*26683^ heterozygotes, were infected intraperitoneally with 5×10^7^ *M. leprae* isolated from mouse footpads; bacteria were tested for viability by radiorespirometry, as described [1]. *rag1*^**t*26683/*t*26683*^ and *rag1*^*+/*t*26683*^ were identified among offspring from a *rag1*^*+/*t*26683*^ incross by genotyping using high-resolution melt analysis of amplicons generated with primers GCGCTATGAGATCTGGAGGA and TGCAGTGCATCCAGAGTAGG, or GCGCTATGAGATCTGGAGGA and CAGAGTAGGCTGGGTTTCCA, on a CFX Connect Thermocycler (BioRad). Animals were observed twice daily and killed by tricaine overdose for each experimental time point, or in the survival experiment, if they appeared moribund. Sections were prepared for histology as described [6]. Briefly, serial sagittal sections were made from formalin-fixed animals and stained by hematoxylin and eosin to visualize host cells, and using Fite, a modified acid-fast stain to visualize *M. leprae* which are acid fast bacilli (AFB). Sections were examined using bright field microscopy and images were collected with a digital photo camera (model DKC-5000; Sony, Tokyo, Japan) and produced using Metamorph software (Molecular Devices Corporation, Sunnyvale, CA). Three fish per group per time point were examined. As a surrogate for bacterial burden per fish, Tissue Studio 4.0 (Definiens) was used to identify the AFB-positive regions in a single sagittal section, and measure their cumulative area. Animals were considered to have cleared infection if no AFB were detected in the entire sagittal section. Statistical analyses were performed using Prism (version 5.0a, GraphPad).

## Results

5×10^7^ *M. leprae* were injected into zebrafish, similar to the number of bacteria used to inoculate mouse footpads [1]. Within seven days post infection (dpi) with *M. leprae*, zebrafish had formed organized granulomas throughout the body involving the pancreas, liver, intestine, mesentery, gonad and adipose tissue (figure 1A). The granulomas were comprised centrally of macrophages that had undergone epithelioid transformation (characterized by a high cytoplasm to nucleus ratio), with scattered lymphocytes (characterized by a high nucleus to cytoplasm ratio) aggregating at the periphery (figure 1A). Thus, even from this early stage, they resembled the organized granulomas of human leprosy (figure 1B). Fite staining revealed that similar-sized granulomas within the same fish contained varying numbers of bacteria, possibly reflecting ongoing bacterial killing (figure 1C and D).

**Figure 1.**
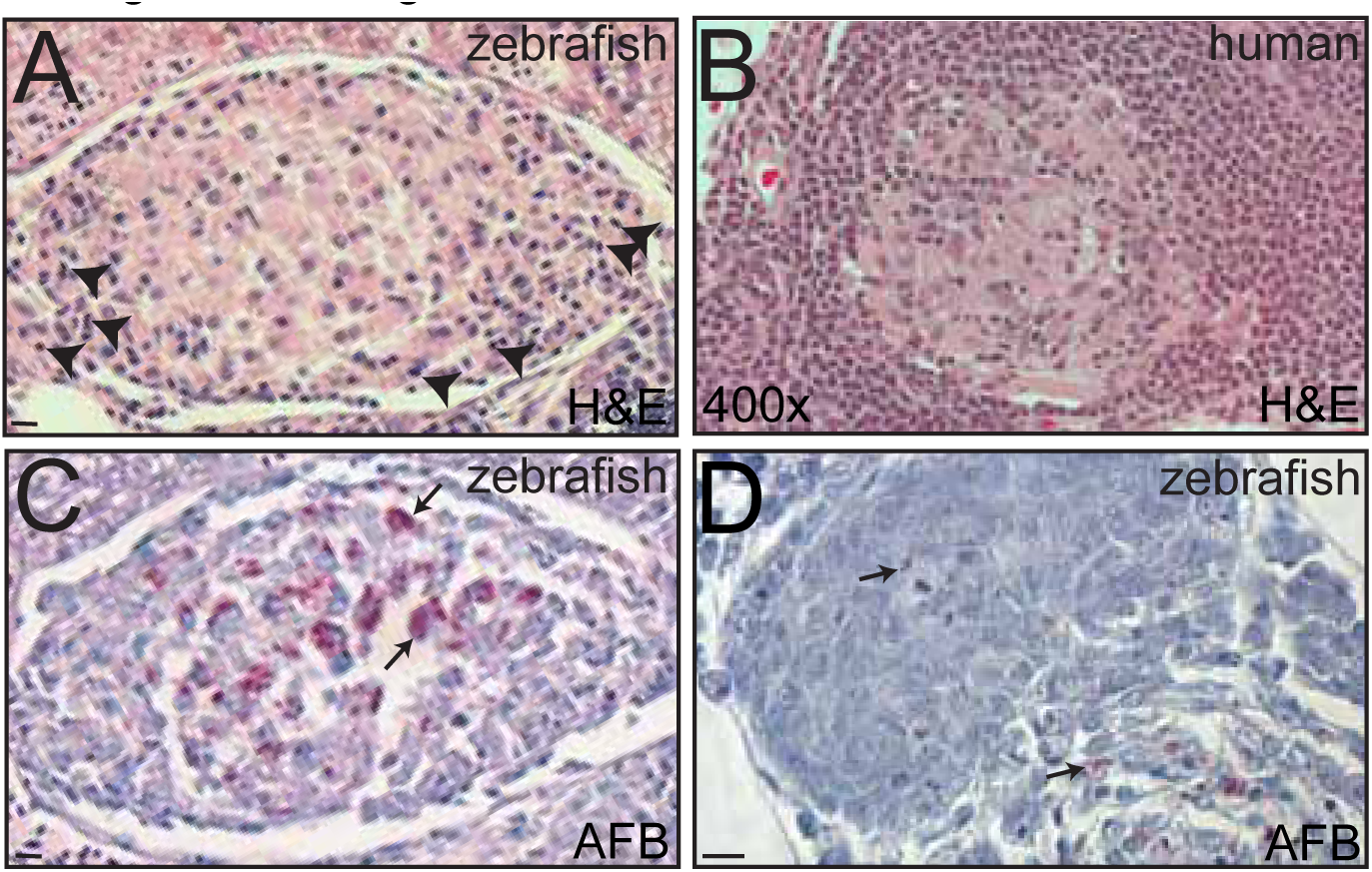
Adult zebrafish are susceptible to *M. leprae* infection. Panel A shows a hematoxylin and eosin (H&E) section of a granuloma in a wildtype adult zebrafish, 7 days post infection (dpi) with 5x10^7^ Thai53 strain *M. leprae*. Arrowheads indicate lymphocyte nuclei. In panel B, a granuloma from a human tuberculoid leprosy patient; Archives of Lauro de Souza Lima Institute. In panel C, a serial section of the granuloma in A, stained for acid-fast bacilli (AFB) to detect *M. leprae*; many bacteria are present (arrows). In panel D, an AFB-stained granuloma section from a similarly infected fish at 7 dpi; few bacteria are present. Arrows indicate bacilli. 10μm bars.

We sought to determine the role of adaptive immunity in the control of leprosy. For tuberculosis, the critical role of adaptive immunity in the control of infection is highlighted by the role of human immunodeficiency virus (HIV) infection in increasing susceptibility to TB [8]. *rag1* mutant mice lacking mature T and B cells are hypersusceptible to *M. tuberculosis* [5]. Likewise, SCID mice, also lacking mature T and B cells, have increased *M. leprae* burdens in their footpads, which decreases upon administration of T cells to the animals [9]. However, the role of adaptive immunity in the control of human leprosy is unclear. On the one hand, lymphocytes are present in the well-organized granulomas of paucibacillary leprosy, similar to the case with human TB granulomas, and an effective cellular response is associated with paucibacillary leprosy [5, 10]. On the other hand, the evidence that HIV infection exacerbates leprosy in humans is scant, with only isolated reports of increased tendency for multibacillary disease, reactions, and relapse [11].

We previously showed that *rag1* mutant zebrafish are more susceptible to *M. marinum*, recapitulating the findings of *rag1* mutant mice infected with *M. tuberculosis* [5, 6]. Therefore, we asked if *rag1* mutant zebrafish were also more susceptible to *M. leprae*. We compared them to their heterozygous siblings, which are as resistant as wildtype fish to *M. marinum* [6]. By ~60 dpi, the infected mutants had become runted with frayed fins (figure 2A) and began to die soon after (figure 2B). Decreased survival was statistically significant in the infected *rag1* mutants but not the other groups (figure 2B), and all dying animals manifested similar signs of disease before death (runting, frayed fins, hemorrhaging, and swimming near the tank bottom). Only 3 of 12 infected mutants survived, and these survivors appeared healthy, suggesting some mutants were able to clear infection.

**Figure 2.**
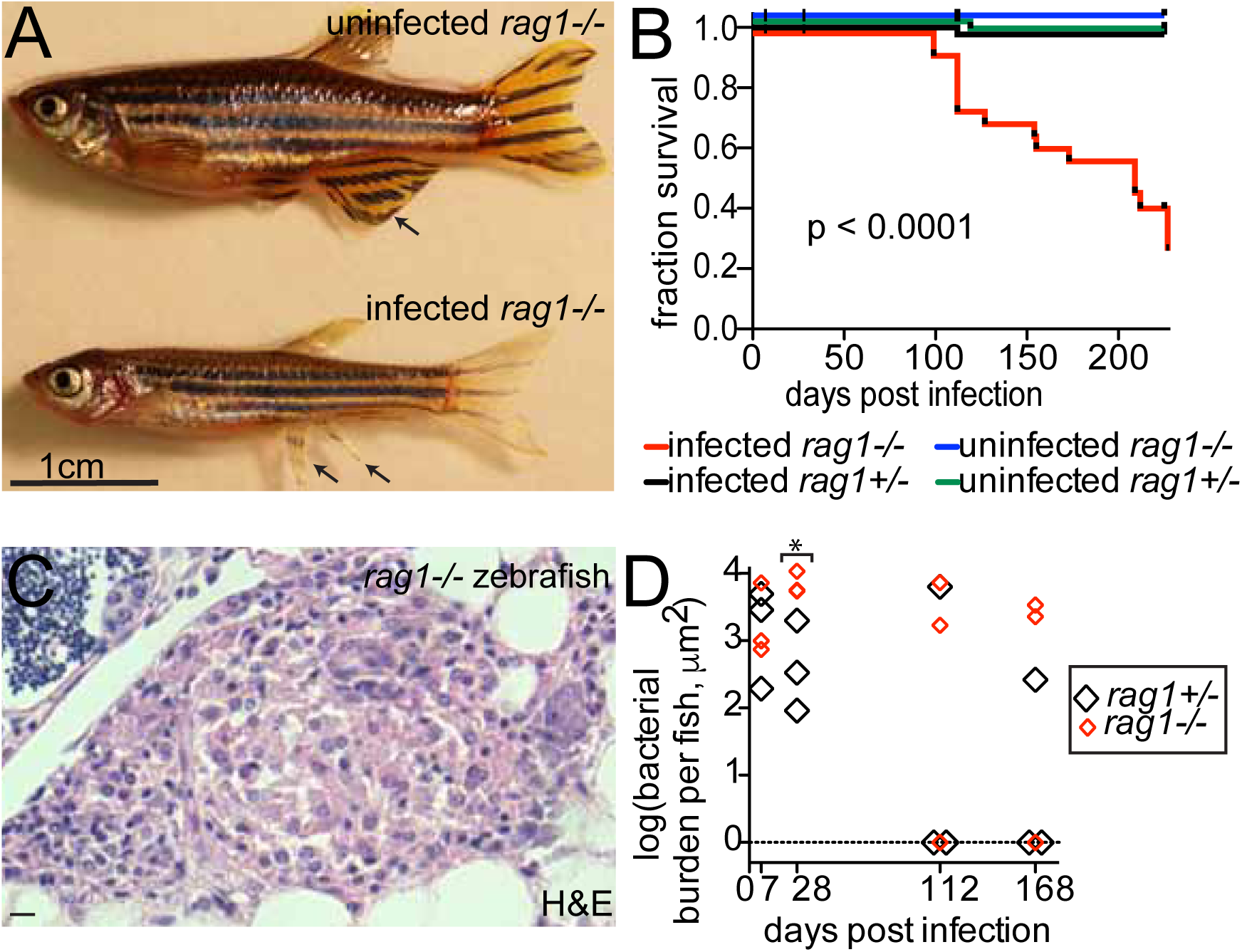
Adaptive immunity contributes to control of *M. leprae* infection. In panel A, representative images of sibling uninfected and infected *rag1* mutant animals ~100 days after infection; the *M. leprae*-infected animal is smaller than the uninfected animal. Arrows indicate an intact fin in the uninfected animal and a frayed fin in the infected animal. In panel B, Kaplan-Meier survival curve of sibling *rag1* heterozygote and mutant zebrafish, infected or not with *M. leprae* as in figure 1A. Number of animals: 61 uninfected heterozygotes, 20 infected heterozygotes, 57 uninfected mutants, 41 infected mutants. In panel C, an H&E-stained section through a *rag1* mutant zebrafish granuloma, infected as in figure 1A; 10μm bar. In panel D, quantification of bacterial burden per fish in *rag1* heterozygotes and mutants; *p=0.03, other comparisons not significant; student’s T test comparing heterozygotes to mutants at each time point.

Simultaneously, in a separate small cohort (three *rag1* heterozygote and three mutant animals per time point), we performed tissue histology to assess granuloma morphology and bacterial burdens. *rag1* mutants formed organized epithelioid granulomas by seven days that were similar to wildtype except that, as expected, they lacked lymphocytes (figures 2C). Analysis of Fite-stained histology sections suggested that both heterozygotes and mutants cleared infection over time. At 112 dpi and 168 dpi, two of three *rag1* heterozygotes contained no bacilli, while one of three *rag1* mutants contained no bacilli at those time points (figure 2D). Quantification of bacterial burdens in the remaining fish showed that mutant bacterial burdens were greater than heterozygotes at 28 days but then declined (figure 2D). Together, these findings suggest that *M. leprae* can be controlled by zebrafish without adaptive immunity.

A curious feature of *M. leprae* granulomas is that they seldom become necrotic, even when laden with organisms [10]; this is in sharp contrast to human tuberculous granulomas [5]. In the zebrafish too, we found that even multibacillary lesions where individual macrophages were packed with bacteria seldom became necrotic (figure S1A). Necrosis was observed in only 2.9% of heterozygote granulomas (1 of 34 granulomas in 12 animals) (figure S1B). Similarly, only a minority of the *rag1* mutant granulomas became necrotic – 14%, or 7 of 50 granulomas in 12 animals; this difference was not statistically significant.

Finally, human leprosy granulomas are frequently associated with damage to peripheral nerves. We were unable to assess nerve damage in this study, as even an experienced neuropathologist was unable to identify the nerves in these small animals. In a companion study using zebrafish larvae, which are transparent, we have been able to show the association between early macrophage aggregates and nerve injury (Madigan et al., submitted).

## Discussion

This pilot study already suggests that the adult zebrafish will be an excellent model for studying *M. leprae* granuloma formation and function, and the immune pathways that determine host susceptibility to leprosy. Morphologically, the granulomas resemble those of paucibacillary (or tuberculoid) human leprosy, and like their human counterparts, they are effective in controlling infection [13]. Indeed, the vast majority of humans appear to clear *M. leprae* infection [13], and the zebrafish do as well.

Another intriguing feature of human leprosy is the rarity of granuloma necrosis [10], and this too is preserved in the zebrafish. This could be because *M. leprae* has lost determinants present in *M. marinum* and *M. tuberculosis* that promote granuloma macrophage necrosis.

Finally, our work reveals the complexity of the interplay between innate and adaptive immunity in the control of leprosy. In separate work, we have developed the larval zebrafish as a leprosy model, and we find that macrophages can aggregate into granulomas and control *M. leprae* to a substantial extent in the sole context of innate immunity (Madigan et al., submitted). Our findings here with the *rag1* mutant reinforce the idea that bona fide epithelioid granulomas form without adaptive immunity [5], yet the full microbicidal capacity of the granuloma macrophages requires stimulation by adaptive immunity. Indeed, we find that lymphocytes begin to arrive in the granuloma by seven days after infection, and bacterial burdens diverge between *rag1* heterozygotes and mutants by 28 days (figure 2D). Thereafter, bacterial burdens drop even in the *rag1* mutant fish, suggesting that innate immune factors can gradually control infection (figure 2D). The finding that mutants slowly reduce bacterial burdens, and occasionally even clear infection, suggests that innate immunity alone may be sufficient to control this slowly growing pathogen. The decreased survival of *rag1* mutants in the face of this delayed control may reflect the adverse consequences of chronic infection, or be due to cytokine dysregulation in the absence of adaptive immunity. In any case, our zebrafish findings may reflect the lack of an obvious link between exacerbation of leprosy and HIV co-infection [11]. Moreover, given that innate immunity has a role in clearing infection, the development in humans of multibacillary rather than paucibacillary leprosy may well reflect innate immune deficiencies, some of which are beginning to be identified [10, 15]. It is our hope that these can be broadly identified and studied in the zebrafish, using the publicly available libraries of zebrafish mutants have been generated by chemical mutagenesis and CRISPR technologies [16].

## Acknowledgements

Live *M. leprae* was provided by the HHS/HRSA/HSB/National Hansen’s Disease Program, Baton Rouge, LA., with financial support from NIAID IAA-2646. We thank Christine Cosma for advice on and technical assistance with zebrafish infections, Paul Edelstein for discussions, advice on histology preparations and statistics, Robert Modlin for advice on histology analysis, Philip Scumpia for advice on histology interpretation, and Paul Edelstein and Robert Modlin for manuscript review. This work was supported by an NIH T32 AI1007411 and an NIH NRSA postdoctoral fellowship AI104240 (C.A.M.), and NIH R37AI054503 and the NIH Director’s Pioneer Award (L.R.). C.A.M. is an A.P. Giannini Foundation Postdoctoral Fellow and L.R. is a Wellcome Trust Principal Research Fellow.

## Conflict of interest statement

all authors declare no conflict of interest.

---

**Supplemental Figure 1.** Few *M. leprae* granulomas undergo necrosis. In panel A, an AFB-stained section of a non-necrotizing granuloma in a *rag1* heterozygote zebrafish with heavily infected macrophages (arrows). In panel B, AFB and H&E sections of a necrotic granuloma observed in *M. leprae*-infected *rag1* heterozygote fish. 10μm bars.

## References

1. Truman, R.W. and J.L. Krahenbuhl, Viable M. leprae as a research reagent. International Journal of Leprosy and Other Mycobacterial Diseases, 2001. 69(1): p. 1–12.

2. Job, C., et al., Electron Microscopic Appearance of Lepromatous Foodpads of Nude Mice. International Journal of Leprosy and Other Mycobacterial Diseases, 2003. 71(3): p. 231–239.

3. Sharma, R., et al., The armadillo: a model for the neuropathy of leprosy and potentially other neurodegenerative diseases. Disease Models and Mechanisms, 2013. 6(1): p. 19–24.

4. Wang, H., et al., An in vitro model of Mycobacterium lepraeinduced granuloma formation. BMC Infectious Diseases, 2013. 13(1): p. 279.

5. Ramakrishnan, L., Revisiting the role of the granuloma in tuberculosis. Nat Rev Immunol, 2012. 12(5): p. 352–366.

6. Swaim, L.E., et al.., Mycobacterium marinum Infection of Adult Zebrafish Causes Caseating Granulomatous Tuberculosis and Is Moderated by Adaptive Immunity. Infection and Immunity, 2006. 74(11): p. 6108–6117.

7. Couret, M., The behavior of bacillus leprae in coldblooded animals. Journal of Experimental Medicine, 1911. 13: p. 576–589.

8. Kwan, C.K. and J.D. Ernst, HIV and Tuberculosis: a Deadly Human Syndemic. Clinical Microbiology Reviews, 2011. 24(2): p. 351–376.

9. Azouaou, N., et al., Reconstitution of Mycobacterium leprae immunity in severe combined immunodeficient mice using a T-cell line. International Journal of Leprosy and Other Mycobacterial Diseases, 1993. 61(3): p. 398–405.

10. Renault, C.A. and J.D. Ernst, Mycobacterium leprae [Leprosy], in Mandell, Douglas, and Bennett’s Infectious Disease Essentials, J.E. Bennett, R. Dolin, and M.J. Blaser, Editors. 2017, Elsevier: Philadelphia, PA. p. 371.

11. Lockwood, D.N.J. and S.M. Lambert, Human Immunodeficiency Virus and Leprosy: An Update. Dermatologic Clinics, 2011. 29(1): p. 125–128.

12. Madigan, C., et al., A macrophage response to Mycobacterium leprae phenolic glycolipid initiates nerve damage in leprosy. submitted.

13. Lara, C. and J. Nolasco, Self-healing, or abortive, and residual forms of childhood leprosy and their probable significance. International Journal of Leprosy and Other Mycobacterial Diseases, 1956. 24(3): p. 245–263.

14. Yang, D., et al., Mycobacterium leprae-Infected Macrophages Preferentially Primed Regulatory T Cell Responses and Was Associated with Lepromatous Leprosy. PLOS Neglected Tropical Diseases, 2016. 10(1): p. e0004335.

15. Tobin, D.M., et al., The lta4h Locus Modulates Susceptibility to Mycobacterial Infection in Zebrafish and Humans. Cell. 140(5): p. 717–730.

16. Varshney, G.K., R. Sood, and S.M. Burgess, Chapter One - Understanding and Editing the Zebrafish Genome, in Advances in Genetics, J.C.D. Theodore Friedmann and F.G. Stephen, Editors. 2015, Academic Press. p. 1–52.

